# Identification and characterization of neoantigen-reactive CD8+ T cells following checkpoint blockade therapy in a pan-cancer setting

**DOI:** 10.1101/2024.03.17.585416

**Authors:** Keith Henry Moss, Ulla Kring Hansen, Vinicius Araújo Barosa de Lima, Annie Borch, Esteban Sanchez Marquez, Anne-Mette Bjerregaard, Østrup Olga, Amalie Kai Bentzen, Andrea Marion Marquard, Mohammed kadivar, Inge Marie Svane, Ulrik Lassen, Sine Reker Hadrup

**Author notes:** These authors contributed equally (Keith Henry Moss, Ulla Kring Hansen, Vinicius Araújo Barosa de Lima, Annie Borch).

## Abstract

**Background:** Immune checkpoint blockade (ICB) has been approved as first-line or second-line therapies for an expanding list of malignancies. T cells recognizing mutation-derived neoantigens are hypothesized to play a major role in tumor elimination. However, the dynamics and characteristics of such neoantigen-reactive T cells (NARTs) in the context of ICB are still limitedly understood.

**Methods:** To explore this, tumor biopsies and peripheral blood were obtained pre- and post-treatment from 20 patients with solid metastatic tumors, in a Phase I basket trial. From whole-exome sequencing and RNA-seq data, patient-specific libraries of neopeptides were predicted and screened with DNA barcode-labeled MHC multimers for CD8^+^ T cell reactivity, in conjunction with the evaluation of T cell phenotype.

**Results:** We were able to detect NARTs in the peripheral blood and tumor biopsies for the majority of the patients; however, we did not observe any significant difference between the disease control and progressive disease patient groups, in terms of the breadth and magnitude of the detected NARTs. We also observed that the hydrophobicity of the peptide played a role in defining neopeptides resulting in NARTs response. A trend towards a treatment-induced phenotype signature was observed in the NARTs post-treatment, with the appearance of Ki67^+^ CD27^+^ PD-1^+^ subsets in the PBMCs and CD39^+^ Ki67^+^ TCF-1^+^ subsets in the TILs. Finally, the estimation of T cells from RNAseq was increasing post versus pre-treatment for disease control patients.

**Conclusion:** Our data demonstrates the possibility of monitoring the characteristics of NARTs from tumor biopsies and peripheral blood, and that such characteristics could potentially be incorporated with other immune predictors to understand further the complexity governing clinical success for ICB therapy.

## Introduction

Early preclinical studies have suggested evidence for the PD-1/PD-L1 axis in tumor-driven immune suppression and the activation of this signaling pathway has resulted in the evasion of antigen-specific T cell responses (1,2). Phase I studies initially investigated monoclonal antibodies targeting PD-1 and PD-L1 in advanced solid tumors, which led to the development of the first checkpoint inhibitors, Nivolumab and Pembrolizumab, and approval by the FDA (1,2). The initial approval of Pembrolizumab (α-PD-1) for advanced or unresectable melanoma set the scene for immunotherapy and moved it to center stage(3). Accordingly, antibodies targeting the PD-1/PD-L1 pathway have been approved as first-line or second-line therapies for an ever-expanding list of malignancies, including lung cancers, renal cell carcinoma (RCC), lymphoma, head and neck squamous cells cancer (HNSCC), bladder cancer and gastro-esophageal cancer. However, these forms of therapies are not effective for the majority of patients receiving immune checkpoint blockade (ICB), despite major advancements in the field. As a result, there is an interest to understand the key parameters that dictate the outcome of therapy, in the setting of ICB.

Among these parameters is the tumor mutational burden (TMB). TMB and neoepitopes have been shown to predict clinical response to immunotherapies based on checkpoint inhibition (4). However, in other studies these have also been shown to be of weak predictive value (5). Recently proposed cancer immune predictor suggests using a combination of biomarkers to better predict the response of ICB (6,7). The neoepitope load is based on predicted neoepitopes, however, with only 2-4% of these epitopes giving a T cell response, studies have tried to explore for more precise prediction strategies (8) or to validate the presence of truly immunogenic neoepitopes as a more well-suited immune predictor (9,10). Additionally, CD39 expression on CD8+ T-cells has been discovered as a marker to distinguish neoepitope-specific CD8+ T cells from bystander CD8+ T cells (11) and more recently, a molecular signature characterizing the neoantigen-specific CD8+ T-cells has been suggested (12) Potentially shifting the focus from TMB as an immune predictor, towards profiling T cell state and function in the context of neoantigen-specific CD8+ T cells in ICB.

In this study, we use DNA barcode-labeled MHC multimers to detect neoantigen-reactive CD8^+^ T cells (NARTs) in clinical samples from a Phase I basket trial. We monitor the dynamics of NARTs and their phenotypic properties along the course of ICB therapy in multiple cell compartments to investigate the hypothesis that the presence of NARTs, and their characteristics, can be used as an immune predictor for clinical outcome to ICB therapy in a pan-cancer setting.

## Results

### Neoantigen-reactive CD8^+^ T cells are detectable in a diverse cohort of cancer patients

T cell recognition toward neoantigens was evaluated in a basket-trial cohort, consisting of 20 cancer patients diagnosed with nine different cancer types and treated with seven different ICB therapies targeting the PD-1/PD-L1 axis, either as a mono or combination therapy. Clinical response to treatment was assessed according to the response evaluation criteria in solid tumors (RECIST) version 1.1, where the best-obtained response, lasting for at least 2 months, was reported. Based on these criteria, the patients were grouped into non-progressive disease (CR, PR, SD. n = 7, and progressive disease (PD. n = 13). Clinical information is presented in Supplementary Table 1.

The experimental workflow for the detection and phenotypic analysis of NARTs is schematically depicted in Figure 1A. Patient-specific neopeptide were predicted with mutant peptide extractor and informer (MuPeXi) (13) from whole exome sequencing (WES) and RNA sequencing (RNAseq). Neopeptide libraries were then synthesized and assembled with DNA barcode-labeled pMHC multimers into patient-specific multimer panels. In total 5383 unique pMHC have been screened, on average 270 per patient covering 1 to 4 different alleles per patient including HLA-A and HLA-B as HLA-C alleles were excluded due to non-TCR driven binding to pMHC. This resulted in 166 NARTs in total with an average of 8 per patient. Included in each library were HLA-matching epitopes derived from common viruses; cytomegalovirus (CMV), Epstein-Barr virus (EBV), and influenza virus (FLU), for detection of viral antigen-reactive T cells (VARTs). Patient PBMCs were obtained from pre- and post-therapy and were then stained with the patient-specific multimer panels in conjunction with a 12-parameter T cell phenotype panel. PBMCs from two healthy donors, as controls. Representative flow plots show the gating strategy applied for sorting NART and VART CD8^+^ T cells from bulk PBMCs (Supplementary Figure 1A).

**Figure 1.**
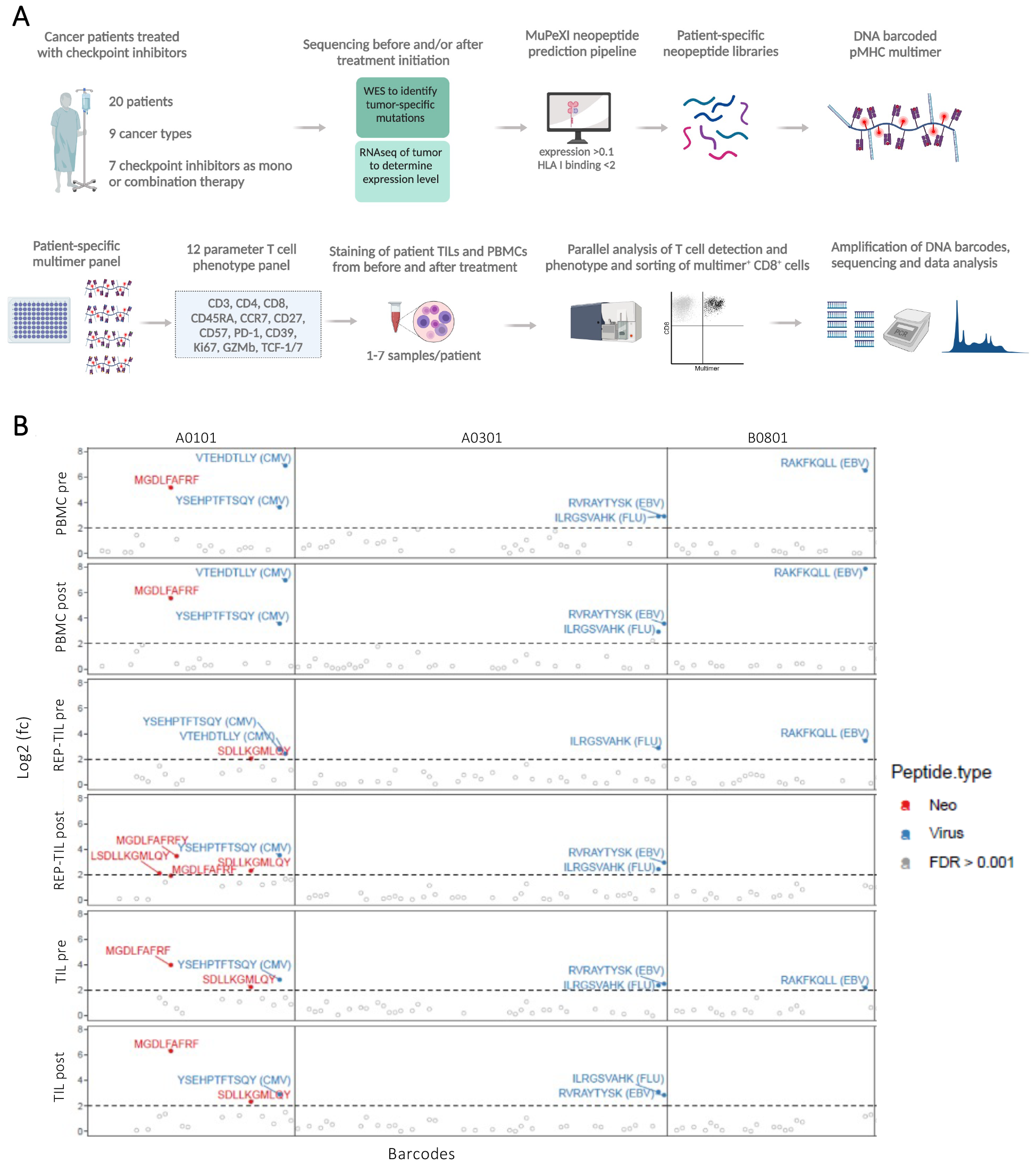
Neoantigen-reactive CD8^+^ T cells are detectable in a diverse cohort of cancer patients. **A)** Experimental workflow for the detection and phenotypic analysis of neoantigen-reactive T cells (NARTs). In total, for all patients, 5383 unique pMHC were included in the screening resulting in 116 NARTs. **B)** Summary plot of the detected T cell reactivity for a representative patient (RH35). This plot is segregated vertically by the different PBMC and TIL samples, and horizontally by the HLA types included in the patient-specific panel. The Log2 fold change (fc) depicts DNA barcodes that have been positively enriched in the given sample, and a threshold Log2 (fc) value of > 2 depicts where significant T cell reactivity (p < 0.001) has been detected towards the neopeptide (red) or viral control peptide (blue).

For each patient, a number of different samples (both TIL and PBMC derived) was evaluated for T cell recognition of neoepitopes. Depicted in Figure 1B is a summary plot of the T cell recognition identified for a representative patient (RH35). This plot is segregated vertically by the different PBMC and TIL samples, and horizontally by the HLA types included in the patient-specific panel. The Log_2_ fold change (fc) depicts enrichment of pMHC associated barcodes in the pMHC multimer positive T cell pool within the given sample, with Log_2_ (fc) > 2 corresponding to significant enrichment (p < 0.001) and presence of a T cell population recognizing the given pMHC, either neoepitopes (red) or viral-derived epitopes (blue). Several NARTs were detected in multiple of the patient’s samples with four unique neoepitopes detected in at least one sample (Figure 1B). T cells recognizing the A0101-restricted neopeptides, MGDLFAFRF, appeared in both the PBMC and TIL samples, and SDLLKGMLQY, appeared in only the TIL and REP-TIL samples. Moreover, corresponding cells recognizing CMV-, EBV-, and FLU-derived epitopes were detected in several of the patient samples.

Two additional patient representative summary plots of T cell recognition is displayed in Supplementary Figure 1B and 1C. For patient 25 (RH25), T cell recognition towards the B2705-restricted neopeptide, KRVFILLLS, was observed in PBMC samples both pre- and post-therapy. Note that this patient has an additional PBMC sample taken from day 86 post-treatment start, and the KRVFILLLS response was also detected there (Supplementary Figure 1B). For patient 27 (RH27), T cell recognition towards the A0201-restricted neopeptide, LLVFLVIYL, was found in both PBMCs and TILs, pre- and post-therapy (Supplementary Figure 1C). Thus, determined by the individual patient’
ss T cell recognition map, variations in T cell recognition can be observed across the different samples, but neoepitope-specific T cells are detectable both pre- and post-therapy in both the blood (PBMC) and tumor (TIL) compartment. Two healthy donor samples were included as controls during the NART screening, and repeatedly used throughout the study. To validate the technical robustness of the T cell detection strategy we correlated the detected of virus-specific T cell populations in these healthy donor samples repeatedly (Supplementary Figure 1D and 1E). We find a strong correlation, related to both Log_2_ (fc) and estimated frequency when the same sample is evaluated for T cell recognition towards viral-derived epitopes, and hence demonstrating robustness in the T cell detection strategy across the different pMHC multimer libraries generated throughout this screening effort.

### NART detection and characterization in PBMC and TILs, pre- and post-therapy demonstrates large variability between patients

NARTs have previously been shown to correlate with clinical outcome (9,10). To assess if the presence of NARTs could be used as an immune predictor for clinical outcome to ICB in this diverse cohort, the total number of NART responses detected across all the patients was assessed, in both PBMCs, TILs, and REP TILs. The number of patient samples and detected responses are given in Supplementary Table 2.

Firstly, to provide an overview of the findings from the T cell screening, we gather information on a per-patient basis, related to the number of neoepitopes included for screening, the numbers of detected NART, the clinical outcome based on RECIST, the cancer and treatment type, and the time to progression (PFS) and overall survival (OS) (Figure 2A). Patients are listed based on the no. of NARTs observed, but no immediate separation can be observed related to patient outcome. Out of the 7 patients where no NARTs are observed, five are classified with PD and two with CR. Importantly, these two CR patients were only screened for 1 or 2 HLA alleles, respectively, due to technical limitations, and hence NART response may be present in restriction to the other HLA molecules expressed by the patients. To further investigate any treatment-related effect on the no. of NART, we evaluated separately the presence of the total number of unique NARTs in PBMCs and TILs, pre- and post-therapy. The no. of NART depicted here, reflects the breadth of neoantigen recognition and the sum of estimated frequency reflects the magnitude of NARTs where neither the breath or the magnitude resulted in a significant difference in relation to the patient clinical outcome (Figure 2B and C). We similarly evaluated NART responses in PBMC and TIL compartments independently, where similar observation was made in both pre- or post-therapy (Supplementary Figure 2A and B). However, when comparing the correlation between the number of detected NARTs in TIL and PBMC compartments, a stronger correlation is found post-therapy compared to pre-therapy (Pearson correlation (0.7>0.5) (Figure 2D), which may indicate an effect on the TILs due to treatment.

**Figure 2.**
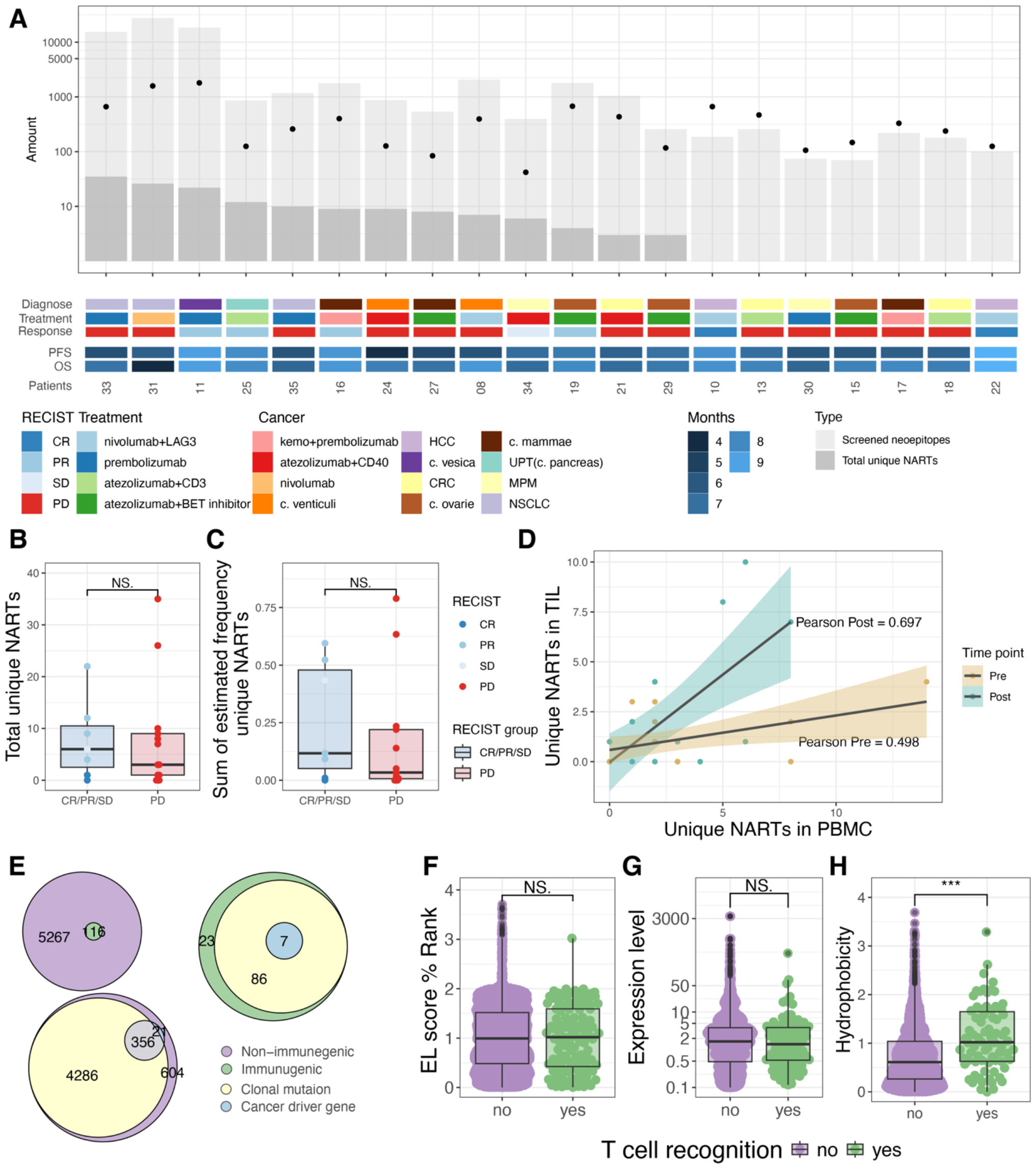
NARTs correlation with outcome and neoepitope immunogenicity. **A)** overview of predicted, screened, and found neoepitopes. **B)** Number of found NARTs per patient comparing outcome (p = 0.69). **C)** Sum of estimated frequency per patients comparing the patient outcome (p = 0.5). **D)** Correlation between number of NARTs in TIL and PBMC for pre- and post-treatment. **E)** The number of clonal mutations and driver mutations in the immunogenic and non-immunogenic pool of peptides. No significant enrichment of either the clonal or driver mutations in immunogenic pool of peptides is found. **F-H)** Comparing immunogenic vs. non-immunogenic neotpitopes according to (F) the MHC-binding in Eluted Ligand % Rank (p = 0.85). (G) Expression level of the corresponding gene with where the mutation is found (p = 0.2). (H) The hydrophobicity of the peptide (p=5.6 · 10^−7^).

To assess the kinetics of the detected NARTs between the treatment time points, we estimated the frequencies of NARTs out of CD8^+^ T cells, i.e. magnitude of NARTs, in both PBMCs (Supplementary Figure 2C) and TILs (Supplementary Figure 2D). The estimated frequency is an approximation based on the fraction of read counts for a given neoepitope, obtained from the barcode sequencing, and the frequency of multimer^+^ CD8 T cells assessed during cell sorting.

Data points represent the sum of estimated frequencies for each individual patient, i.e. the total magnitude to the NART responses per patient. In the PBMCs, for both patient groups, there is a tendency that the magnitude of the detected NARTs decreases at day 34 (1st time-point, post therapy). In the TILs, mixed tendencies are observed, where the magnitude of NARTs for some patients increases post-therapy, and for others, declines.

In general terms, the number of NARTs detected was variable among patients, with most patients exhibiting very limited neoepitope T cell recognition. Also, no difference was observed in the number of predicted neoepitopes, nor the TMB across this patient cohort, when compared related to clinical outcome and pre-post therapy effect (Supplementary Figure 2E and F). Interestingly, although the TMB, predicted neoepitopes, and detected NART remains relatively stable over the course of treatment, we do observe that patients with non-progressive seem to have more homogeneous tumor characteristics than progressive disease patients (Supplementary Figure 2G+H), with an overlap between pre- and post-tumor biopsies in regard to non-synonymous mutations and predicted neoepitopes being 27% and 24%, among non-progressive disease patients, and 19% and 15%, respectively, among progressive disease patients. The lack of overlap can also be related to technical variability and tumor heterogeneity, but as all patient samples are collected and processed similarly, the difference observed between the non-progressive and progressive disease group can be attained to their mutational characteristics.

To characterize additional aspects related to CD8^+^ T cell recognition of neoepitopes, we determined if the gene the mutation arrives from is related to a cancer driver gene or found clonally. Here no difference was found when comparing the immunogenic to the non-immunogenic pool of neoepitope candidates (proportion z-test) (Figure 2E). We assessed Eluted ligand (EL) rank score, expression level, and hydrophobicity across all patients to explore these characteristics in regards to neoepitope immunogenicity. No difference was found when observing the MHC-I binding (EL%Rank) (Figure 2F) or the expression level (Figure 2G). Of note, both the expression and pMHC binding were used for the pre-selection of the neoepitope candidates included for T cell recognition screening, which could explain why no further influence was observed for thede parameter. Interestingly we observed the peptide hydrophobicity to be significant different in the immunogenic versus non-immunogenic fractions of the neoepitopes (Figure 2H, p = 5.6x10^−7^). With stronger hydrophobicity increasing the chance for T cell recognition of the given neopeptide.

Thus, taken together, NART populations can be detected in 13 of 20 patients in this diverse cohort, often both in PBMC and TIL. NART responses were observed both pre- and post-therapy. No influence on treatment outcome could be found, but a trend toward an increasing magnitude of NART recognition was observed in non-progressive disease patients. When investigating the neoepitope characteristics, hydrophobicity was identified as a relevant feature to potentially include for neoepitope prediction for the future.

### Phenotypic differences of bystander CD8^+^ T cells and NARTs

To comprehensively evaluate T cell characteristics, a multiparametric analysis was conducted using the dimensionality reduction technique, UMAP, in combination with unsupervised clustering by FlowSOM. The global structure of the bulk CD8^+^ T cells and NARTs, from all patient PBMCs, without grouping, is depicted as density plots in Figure 3A. Overlays of pre-treatment (light blue) versus post-treatment (red) are given alongside the density plots, for both bulk CD8^+^ T cells and NARTs. FlowSOM clustering was subsequently overlaid on the UMAP to generate a profile for the various subpopulations (Figure 3B). When looking at the density plots and overlay for bulk CD8^+^ T cells, with reference to the selected subpopulations (Figure 3C), it was apparent that the black and pink subpopulations seemed to reduce post-treatment, whereas the orange one seemed to increase. This corresponded with the following phenotypic profiles: black (Ki67^hi^, CD27^hi^, GZMb^int^), pink (Ki67^hi^, CD27^low^, GZMb^int^) and orange (CD57^hi^, GZMb^int^). Similarly, when looking at the NARTs specifically, it is apparent that the orange, light green and dark green subpopulations seemed to increase post-therapy. This corresponded with light green (CD57^hi^, CD45RA^hi^, GZMb^hi^) and dark green (CD27^hi^, TCF-1^int^). The plotting of the selected subpopulations does not seem to reveal any significant treatment-induced phenotypic signature, or any profile difference related to either the bulk CD8^+^ T cells or NARTs (Figure 3D).

**Figure 3.**
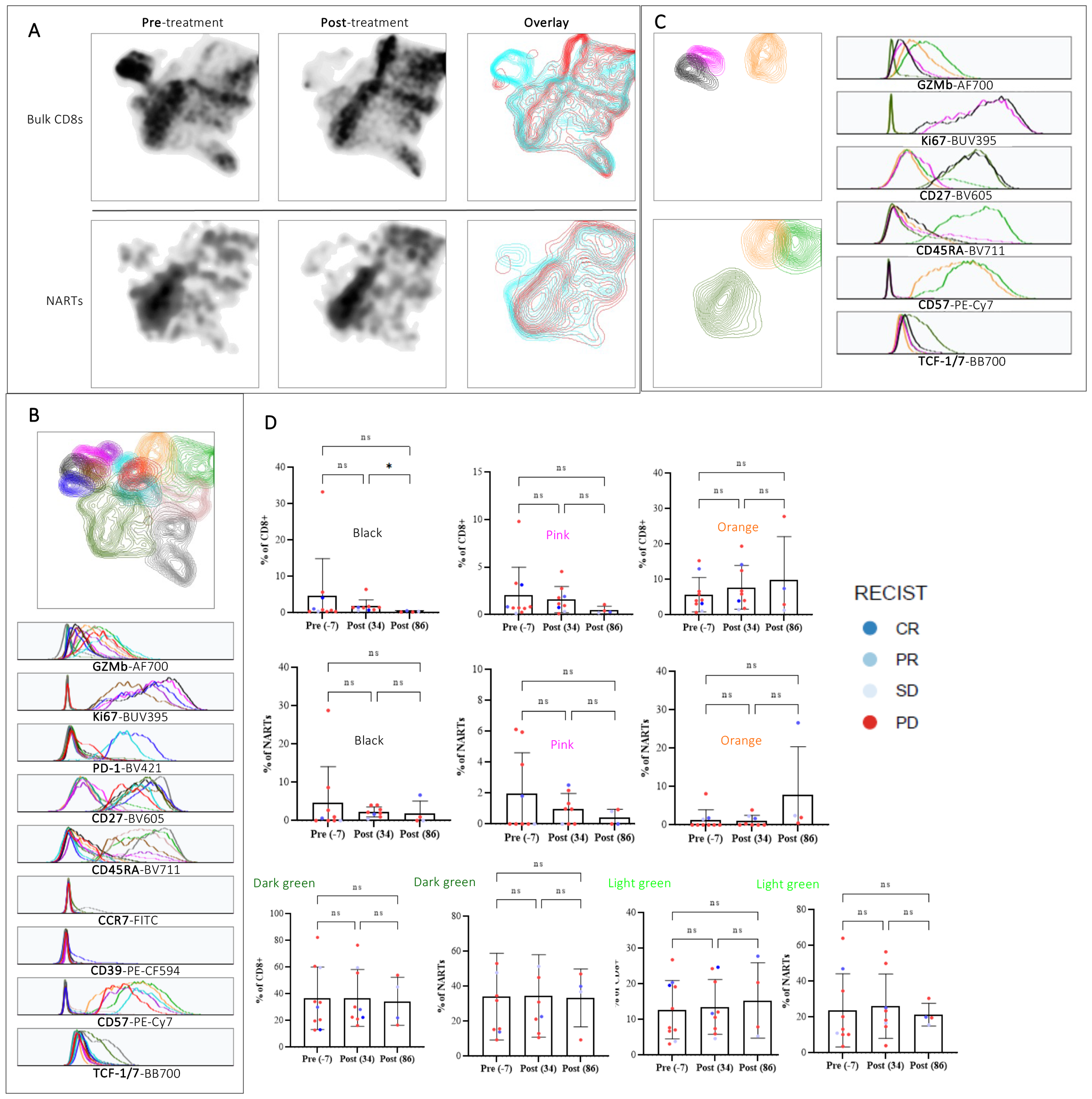
Phenotypic characterization of bulk CD8^+^ T cells and NARTs from patient-derived PBMCs. **A)** UMAP representation of the global structure of bulk CD8^+^ T cells (top) and NARTs (bottom) from all patients, pre- and post-treatment depicted as density and overlay plots (light blue: pre-treatment, red: post-treatment). **B)** Overlay of FlowSOM clustering on UMAP, with conjunct histograms to visualize and identify the sub-populations within the global structure. **C)** Selected sub-populations from the FlowSOM clustering and histograms for the corresponding phenotypic profiles. **D)** Quantitative assessment of the FlowSOM clustering for the selected sub-populations, depicting the frequency of CD8+ T cells or NARTs that the particular sub-populations represent. For all the plots related to frequency of bulk CD8+ T cells: (−7, n = 10; 34, n = 9; 86, n = 4) and for all plots related to frequency of NARTs: (−7, n = 9; 34, n = 8; 86, n = 4). Significance for D is denoted based on Kruskal-Wallis test with Dunn’s correction

When focusing specifically on the density plots of the NARTs, there are apparent changes in the global structure post-treatment, particularly in the case of the disease control patients (Supplementary Figure 3A) with the overlay of the two disease groups, disease control (blue) and progressive disease group (red) (Supplementary Figure 3B). The relevant subpopulations were focused on for phenotypic profiling (Supplementary Figure 3C). These three subpopulations of interest, that were prominent post-treatment in the NARTs from the disease control group, corresponded to the following profiles: dark blue (Ki67^hi^, PD-1^hi^, CD27^hi^), light blue (PD-1^hi^, CD27^hi^, CD57^hi^, GZMb^hi^) and red (PD-1^low^, CD27^int^, CD45RA^int^, CD57^int^, GZMb^hi^) (Supplementary Figure 3D). However, no significant differences were found which also can be reflected by the low number of samples when comparing the two disease groups.

### Gene expression analysis from the tumor microenvironment reveals a signature related to treatment outcome

When analyzing the bulk RNAseq data from the TME, an estimate of T cell abundance was determined using the microenvironment cell populations (MCP) counter. Here the patients are grouped according to RECIST criteria, and the T cell abundancy is compared pre- and post-treatment, where RNAseq data was available. A significantly increased T cell population was found post-compared to pre-treatment in the CR/PR/PD group (Figure 4A). The same tendency, however non-significant, was found in the cytotoxic lymphocyte estimation from MCP-counter (Figure 4B). Interestingly, the no. of NARTs detected and the T cell infiltration estimated from the MCP-counter showed a better correlation post-treatment (Pearson = 0.4) compared to pre-treatment (Pearson = 0.048) (Figure 4C and D).

**Figure 4.**
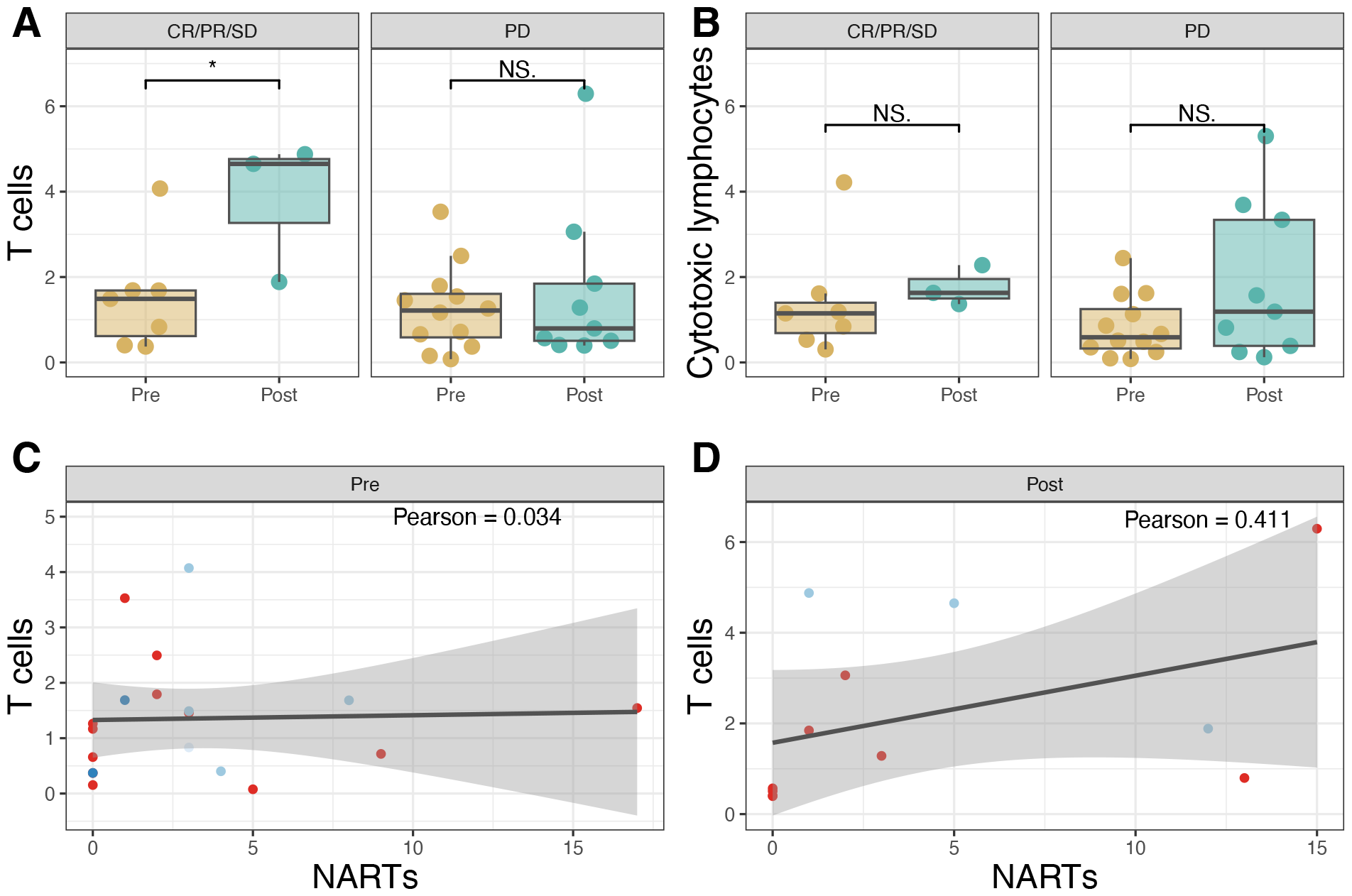
T cell extraction from RNAseq. **A)** T cell estimation from MCP counter. CR/PR/SD (p = 0.033) and PD (p = 0.86). **B)** Cytotoxic lymphocyte estimation from MCP counter CR/PR/SD (p = 0.18) and for PD (p= 0.28). **C+D)** T cell from MCP counter correlation with NARTs before (C) and after therapy (D).

## Discussion

We report herein, a comprehensive evaluation of the characteristics of neoantigen-reactive CD8^+^ T cells, in the context of ICB therapy. While, immune predictors, such as TMB or the presence of inhibitory receptors, provide prognostic value in some patients, there is currently no universal immune predictor for all patients of different cancer types (5,14). Basket trial approaches, where patients are selected for treatment based on their molecular characteristics (e.g. TMB) rather than their diagnosis, are more frequently used in clinical practice. Thus, understanding how immune characteristics may affect treatment outcomes, even in such a diverse cohort as presented here, is of increasing importance.

We initially set out to evaluate whether the characteristics, such as breadth and magnitude of detected NARTs, were able to provide a distinction between the disease control and progressive disease patients. There was no significant difference in the breadth of NARTs between the patient groups, apart from a minor tendency for an increase in the disease control group. In the PBMCs, the breadth of the detected NARTs tended to decrease initially post-treatment, followed by an increase at the later time point, in the disease control group. This seems to confound previous reports showing an early increase in detected NARTs post ICB treatment, e.g. in the case of a bladder cancer study (10). In the progressive disease group, the breadth of detected NARTs tended to decrease and did not show a similar increase at the later time point. There appeared to be no clear tendencies for the breadth of the NARTs from the TILs. Although, recent studies have highlighted that PD-1 axis blockade does not necessarily increase the breadth of the NARTs in the TME(15) and another study showed that only limited immunodominant mutations showed neoantigen-specific immune responses (16). According to the bulk RNAseq data from the TME, the estimated level of T cells significantly increases in post compared to pre-treatment in the disease control group, where no difference was found in the progressive disease group indicating that the response to treatment is related to T cell infiltration. Hence the absence of NARTs in the CR patient could be due to the low HLA-coverage for some patients. The TME also pre-treatment resulted in a separation according to patient outcome, however, no distinct genes were found as the TMB or NARTs did not significantly distinguish the patients based on outcome, as also recent has been described ((6,7) a combination strategy might be more valid at immune predictor, especially for a heterogeneous population.

We next evaluated the magnitude of the NARTs in the PBMCs and TILs, from pre-to post-treatment, to assess for treatment-induced kinetics. There was no significant difference between the groups; however, in a similar fashion to the breadth, there was a tendency that the magnitude of the detected NARTs decreased initially post-treatment followed by an increase at the later time point, at least in the disease control group. This would support a previous finding, where a higher frequency of neoantigen-specific T cells was detected in NSCLC patients responding to anti-PD-L1 therapy(17). However, other reports has suggested that PD-1 blockade provides therapeutic benefit not by increasing T cell infiltration, but rather by reducing the generation of anergic tumor-reactive T cell populations within the TME(15). In the TILs, again, no clear tendencies were observed. Strikingly, only 18% of the NARTs were detected in both TILs and PBMCs in our study. However, this is consistent with previous findings in an ovarian cancer cohort, where a similar discordance of neoepitope recognition was observed between TILs and PBMCs (18). This discrepance can also relate to a lower frequency of NARTs in circulation, and hence challenge their detection.

The characteristics of neoepitope immunogenicity revealed that neither the predicted MHC binding affinity nor the neoepitope expression level was able to predict T cell recognition, within the range tested, but the neopeptide hydrophobicity seemed to influence the neoepitope immunogenicity. Neopeptide hydrophobicity has been recently reported to play a role in T cell receptor (TCR) recognition (19). However, to improve the accuracy of neopeptide prediction, these parameters, and others, may need to be incorporated into a combined model. This is particularly relevant, considering the low fraction of predicted neopeptides that result in T cell recognition ((20).

Considering the reports that the functional state of tumor-reactive T cells may ultimately dictate the therapeutic outcome to ICB (21), we evaluated the phenotypic signature associated with NARTs and bulk CD8^+^ T cells. In the PBMC compartment, a treatment-induced signature was observed for both bulk CD8^+^ T cells and NARTs. Post-treatment, the bulk CD8^+^ T cells seemed to lose two proliferating subsets, of which, one was in an early-differentiated state and the other a late-differentiated state i.e. black (Ki67^hi^, CD27^hi^) and pink (Ki67^hi^, CD27^low^). A late-differentiated effector subset, i.e. orange (CD57^hi^, GZMb^int^), became more prominent. Since these are frequencies, however, this may be due to the loss of other subsets. Two additional subsets were highlighted, which became more prominent in the NARTs. A late-differentiated T_EMRA_ subset i.e. light green (CD57^hi^, CD45RA^hi^, GZMb^hi^) and an early-differentiated subset i.e. dark green (CD27^hi^, TCF-1^int^) appeared to be more prominent post-treatment. Again, since these subsets appear not to be proliferating (Ki67^neg^), this could be due to the disappearance of other subsets. However, T_EMRA_s in the blood have been shown to increase in frequency in the context of prolonged viral infection, but also with age(22,23). Recently a neoantigen CD8^+^ T cell signature (NeoTCR signature) has been determined from single-cell dataset ((12,24) however, in our dataset no specific phenotype signature that could distinguish the NARTs from bulk CD8^+^ T cells.

In the TILs, two CD39^+^ subpopulations became more apparent post-treatment, specifically in the NARTs. One proliferating subset, i.e. red (Ki67^hi^, CD27^int^, CD39^hi^, TCF-1^int^) and another low-proliferating subset, i.e. beige (Ki67^int^, CD27^int^, CD39^hi^, TCF-1^int^). Notably, these two subsets appeared to have increased TCF-1 expression. Both CD39 and TCF-1 have been recently implicated in relation to NARTs and ICB. The expression of CD39 has generally been associated with tumor-reactive T cells, allowing the distinction from bystander T cells in the TME, although, CD39 has generally been linked to a dysfunctional, terminally differentiated phenotype(25–27). TCF-1, the stem cell-like marker, has been recently reported to provide T cells with self-renewal properties, although this tends to get downregulated in terminally exhausted T cells, i.e. in the context of chronic viral infection and cancer (28–31).

## Materials and Methods

### Patient cohort

Twenty patients with solid metastatic tumors referred to treatment with checkpoint inhibitors at the Phase 1 Unit - Rigshospitalet, Copenhagen, Denmark, were included in this study. Inclusion criteria for immune therapy: performance status (ECOG scale) up to 1, age above 18 years, and at least one metastatic lesion accessible for radiologically-guided biopsy. Moreover, patients were required to have at least one measurable lesion according to response evaluation criteria in solid tumors version 1.1 (RECIST v.1.1), which was also used for assessing treatment response. The best-obtained RECIST response lasting for at least 2 months was registered. Clinical data are presented in Supplementary Table 1.

### Patient and healthy donor samples

Fresh tumor biopsies were obtained before initiation of therapy and approximately 40 days after treatment commencement from the same metastatic site. Immediately after collection, the tumor biopsies were stored in media containing RPMI1640 (Thermo Fischer Scientific), penicillin, streptomycin, and fungizone (Bristol-Meyers Squibb) for transport (elapsed time of one hour). Biopsies were minced manually under sterile conditions into smaller fragments and placed into 24-well plates (Nunc, Roskilde, Denmark) containing 2 mL of culture medium (90% RPMI1640, penicillin, streptomycin, fungizone, 10% heat-inactivated Human AB Serum) and 6000 UI/mL IL-2 (Proleukin, Novartis, Basel, Switzerland). Plates were kept in incubators (humidified atmosphere, temperature around 37 °C and 5% CO_2_) for five days. Afterward, half of the medium in each well was removed and replenished with 1 mL of culture medium containing interleukin 2 (same concentration as described above). The same procedure was repeated afterwards every second day. Young TILs were pooled around five-six weeks after the biopsy date and cryopreserved similarly as for PBMCs. Further expansion of REP-TILs *in vitro* was done by irradiated allogeneic feeder cells (PBMCs from healthy donors), anti-CD3 antibody, and IL-2 for 14 days as described by Andersen and colleagues(32). It is important to note that for all of the plotting and statistical analysis of this study, the young TILs and REP-TILs were grouped. This was due to inconsistent TIL/REP-TIL availability among the patients. Additionally, there were no follow-up biopsies taken at day 86, so the TILs represent just two time points. Blood samples were collected for isolation of PBMCs on the same day as biopsies carried out. The third collection of blood samples took place approximately 26 weeks after treatment start if patients were still in treatment. PBMCs were isolated from whole blood by density centrifugation on Lymphoprep (Axis-Shield PoC) in Leucosep tubes (Greiner Bio-One) and cryopreservation in inactivated Human AB serum with 10% DMSO. Samples were initially stored at -80°C alcohol-free freezing containers (Cool cell, Biocision) for 24 hours and then stored at -140°C until further use.

Healthy donor blood samples were obtained from the blood bank at Rigshospitalet, Copenhagen, Denmark. PBMCs were isolated from whole blood as described for the PBMCs from the patients and cryopreserved at -150°C in fetal calf serum (FCS, Gibco) with 10% dimethyl sulfoxide (DMSO, Sigma-Aldrich). All healthy donor and patient materials were collected under approval by the Scientific Ethics Committee of the Capital Region, Denmark, with written informed consent obtained according to the Declaration of Helsinki.

### Molecular analysis of tissue biopsies

Fragments of tumor biopsies stored in RNAlater (Sigma-Aldrich) for DNA and RNA extraction. In short, DNA and RNA were isolated using the AllPrep DNA/RNA kit (Qiagen). For blood samples, genomic DNA was extracted using a Tecan automation workstation (Promega). DNA whole-exome sequencing (WES, Illumina platform) and mRNA expression arrays (Human U133 Plus2.0, Affymetrix) were performed on extracted material.

### Next-generation sequencing data analysis

The WES and RNAseq data were processed according to the Genome analysis tool kit (GATK) best practice guidelines for somatic variant calling(33). First, raw reads were trimmed to a minimum length of 50 bp and quality-checked to a Phred score of 20 using Trim Galore 0.4.0(34), combined with Cutadapt(35) and FastQC(36). The human genome (GCh38) was used to align the reads using the Burrows-Wheeler Aligner(37) version 0.7.16a. Duplicate reads were marked with Picard-tools MarkDuplicates version 2.9.1. Base recalibration and contamination tables was performed with GATK version 4.0.1.1 and used as input for MuTect2(38), with matched tumor and normal samples. RNAseq was processed by TrimGalore (0.4.0) and Kallisto version 0.42.1 was used to find expression information. HLA alleles of each patient were determined by first filtering the reads and aligning them to the HLA region using RazerS version 3.4.0(39). Secondly, Optitype 1.2(40) was used to type each patient’s HLA alleles.

### Assessment of TMB and neoepitope load

The total tumor mutational burden (TMB) is the number of non-synonymous mutations from MuTect2 (39). Neopeptides were selected based on an expression above 0.1 TPM and a predicted binding eluted ligand percentile rank (EL %Rank) score below 2.

### Peptides

All selected mutation-derived and virus control peptides were purchased from Pepscan (Pepscan Presto BV, Lelystad, Netherlands) and dissolved to 10 mM in DMSO and stored at -20 °C.

### Tumor Microenvironment analysis

The T cell abundance estimate was found using MCP-counter(41) the Kallisto output from the bulk RNAseq data and as input to ebecht/MCPcounter packages from GitHub in R version 4.0.2.

### MHC monomer production and generation of specific peptide-MHC complexes

The production of MHC monomers was performed as previously described(42,43). In brief, human β_2_ microglobulin (β_2_m) light chain and the heavy chains of the included HLA types were expressed in bacterial Bl21 (DE3) pLysS strain (Novagen, cat#69451) and purified as inclusion bodies. Followed by folding of heavy chain and β2m light chain complexes with a UV-sensitive ligand(44,45), biotinylation with BirA biotin-protein ligase standard reaction kit (Avidity, 318 LLC-Aurora, Colorado), and purification using size-exclusion column (Waters, BioSuite125, 13µm SEC 21.5 × 300 mm) HPLC (Waters 2489). Specific peptide-MHC (pMHC) complexes were generated by UV-induced peptide exchange(44,46).

### Detection of peptide-MHC specific T cells by DNA barcode-labeled multimers and phenotypic characterization

Patient-specific libraries of predicted neopeptides and virus control peptides were generated as previously described(47). In brief, PE and/or APC - labeled dextran backbones were first coupled with DNA barcodes followed by the pMHC complexes, generated above. Hence, a specific peptide was given a unique DNA barcode together with either a PE or APC-fluorescent label. Patient samples and healthy donor PBMCs were stained with an up-concentrated pool of all multimers in the presence of 50 nM dasatinib. Followed by staining with an antibody mix composed of CD8-BV480 (BD, cat#566121, clone RPA-T8), dump channel antibodies (CD4-FITC (BD, cat#345768), CD14-FITC (BD, cat#345784), CD19-FITC (BD, cat#345776), CD40-FITC (Serotech, cat#MCA1590F), and CD16-FITC (BD, cat#335035)) and a dead cell marker (LIVE/DEAD Fixable Near-IR; Invitrogen, cat#L10119). The samples included for T cell phenotypic characterization were stained with a separate antibody cocktail containing T cell lineage markers (CD3-BV786 (BD, cat. #563799, clone SK7), CD4-BV650 (BD, cat. #563876), and CD8-BV480 (BD, cat. #566121, clone RPA-T8)), characterization markers (Ki67-BUV395 (BD, cat. #564071, clone B56), PD1-BV421 (BD, cat. #562516, clone EH12.1), CD27-BV605 (BioLegend, cat. #302830, clone O323), CD45RA-BV711 (BD, cat. #563733, clone HI100), CCR7-FITC (BioLegend, cat. #353215, clone G043H7), CD39-PE-CF594 (BD, cat. #563678, clone Tu66), CD57-PECy7 (BioLegend, cat. #393310, clone QA17A04), GranzymeB-AlexaFlour700 (BioLegend, cat. #372221, clone 581 QA16A02)), TCF-1/7-BB700 (BD, cat. #353988, clone S33-96C) and a dead cell marker (LIVE/DEAD Fixable Near-IR; Invitrogen, cat. #L10119). Multimer-binding T cells were sorted as lymphocytes, single, live, CD8^+^, FITC^-^ and PE^+^/APC^+^ and pelleted by centrifugation. DNA barcodes were amplified from the isolated cells and a stored aliquot of the multimer pool used for staining (diluted 50,000× in the final PCR reaction, used as a baseline). PCR products were purified with a QIAquick PCR Purification kit (Qiagen, cat#28104) and sequenced using an Ion Torrent PGM 316 or 318 chip (Life Technologies) at PrimBio, USA. Sequencing data were processed by the software package Barracoda (available online https://services.healthtech.dtu.dk/services/Barracoda-1.8/). Barracoda identifies the DNA barcodes annotated for a given experiment, assigns a sample ID and pMHC specificity to each DNA barcode, and counts the total number of reads and clonally reduced reads for each peptide-MHC-associated DNA barcode. A Log_2_ fold change value is estimated through mapping read counts in a given sample relative to the mean read counts of the triplicate baseline samples using normalization factors determined by the trimmed mean of M-values method. FDRs were estimated using the Benjamini–Hochberg method. A threshold of at least 1/1,000 reads associated with a given DNA barcode relative to the total number of DNA barcode reads in that given sample was set to avoid false-positive detection of T cell responses due to a low number of reads in the baseline samples. An estimated cell frequency was calculated for each DNA barcode from the read count fraction out of the percentage of CD8^+^ multimer^+^ T cells. DNA barcodes with a FDR < 0.1%, equal to p < 0.001, were considered to be significant T cell responses. Samples with low viability of CD8^+^ T cells (< 5000 cells) were excluded and DNA barcodes enriched in both patient samples and healthy donor controls were excluded as technical background.

### Confirmatory tetramer staining

Patient-specific neopeptides and viral control peptides were diluted to 200 µM in PBS. MHC monomers were diluted to 150 µg/mL and mixed in a 1:1 ratio (vol) with diluted peptides. pMHC complexes were generated by UV-induced peptide exchange. pMHC complexes were centrifuged at 3300 g, 4C for 5 min and the supernatant was transferred to a new plate. APC-labeled streptavidin (BioLegend, 0.2 mg/mL) was then added stepwise to the pMHC complexes. After a cumulative 30 min of incubation on ice, the MHC tetramers (10ug/mL) were incubated with freshly thawed PBMCs in the presence of 50 nM dasatinib. Followed by staining with an antibody mix composed of CD8-BV480 (BD, cat#566121, clone RPA-T8), dump channel antibodies (CD4-FITC (BD, cat#345768), CD14-FITC (BD, cat#345784), CD19-FITC (BD, cat#345776), CD40-FITC (Serotech, cat#MCA1590F), and CD16-FITC (BD, cat#335035)) and a dead cell marker (LIVE/DEAD Fixable Near-IR; Invitrogen, cat#L10119).

### Flow cytometry and phenotypic analysis

All flow cytometry experiments were carried out on FACSMelody and FACSAria Fusion instruments (BD Biosciences). Data were analyzed in FlowJo version 10.7.1 (TreeStar, Inc). For UMAP dimensionality reduction, PBMCs or TILs were pre-gated on lymphocytes, singlets, live, CD3^+^ and CD8^+^. The CD8^+^ populations were then concatenated for the PBMCs (n = 25) or the TILs (n = 24) and subsequently down-sampled to 200000 cells with DownSample v3.3. UMAP v3.1 was then run with the selection of Ki67, PD-1, CD27, CD57, CCR7, CD45RA, CD39, GZMb and TCF-1, with default settings (Euclidean distance function, nearest neighbor value of 15, and a minimum distance of 0.5). The unsupervised clustering algorithm, FlowSOM v2.9, was then run with the selection of 15 meta clusters.

## Supporting information

Supplementary figures and tabels

## Data analysis

The graphing and statistical analyses were conducted using GraphPad Prism 9 or R statistical software version 4.0.4. The data were assessed for normal distribution using D’Agostino-Pearson normality test. Non-parametric data were analyzed with unpaired Mann-Withney U-test or Kruskal-Wallis test with Dunn’s correction for multiple comparisons. The correlations were analyzed using Pearson’s correlation.

## Acknowledgment

Thanks to Tripti, who constructed all the monomers. Great thanks to all the patients who have participated in the trail.

## Ethics statement

The studies involving human participants were reviewed and approved by Committee on Health Research Ethics for the Capital Region of Copenhagen and the Danish Data Protection Agency. The patients/participants provided their written informed consent to participate in this study.

## Author contribution

KHM: Barcode screening and flow analysis wrote and revised the manuscript. UKH: Barcode screening wrote and revised the manuscript. VL: responsible for taking patient material and revising the manuscript. AB: Predicted neoepitopes and performed bioinformatic analysis, wrote and revised the manuscript ESM: Barcode screening and revised manuscript. AMB: Supervised in the bioinformatics and revised manuscript. OØ: Revised manuscript. AKB: Supervised for barcode technique and revised manuscript. AMM: Bioinformatic support and revised manuscript. MK: Phenotyping support and revised manuscript. IMS: Revised manuscript. UL: Conceived the concept and had patient contact, supported the funding, and revised the manuscript. SRH: Conceived the concept supervised throughout the study, wrote, and revised the manuscript.

