## Supplementary figures and tabels for "Identification and characterization of neoantigen-reactive CD8+ T cells following checkpoint blockade therapy in a pan-cancer setting"

**Supplementary Table 1: Clinical data.** STAD; Stomach adenocarcinoma, LIHC; Liver hepatocellular carcinoma, BLCA; Bladder Urothelial Carcinoma, CRC; Colorectal cancer, HCC; Hepatocellular carcinoma, GAC; Gastric carcinoma, UPT; Unknown primary tumor, BRCA; Breast invasive carcinoma, OV; Ovarian serous cystadenocarcinoma, MPM; Malignant pleural mesothelioma, NSCLC; Non-small-cell lung carcinoma.
[1]Metastatic site.

| Patient ID | Diagnosis | Checkpoint inhibitor treatment | Number of Immuno therapy cycles | RECIST (best obtained) | Biopsy site | Number of previous treatments lines |
|---|---|---|---|---|---|---|
| **RH08** | STAD | Nivolumab + LAG3 antibody | 4 | PD | Liver[1] | 2 |
| **RH10** | LIHC | Nivolumab + LAG3 antibody | 13 | CR | Liver[1] | 2 |
| **RH11** | BLCA | Pembrolizumab | 18 | PR | Lymph node | 1 |
| **RH13** | CRC | Atezolizumab + CD3 agonist | 6 | PD | Liver[1] | 3 |
| **RH15** | OV | Atezolizumab + BET inhibitor | 4 | PD | Subcutaneous | 6 |
| **RH16** | BRCA | Chemo + Pembrolizumab | 11 | PR | Lymph node | 3 |
| **RH17** | BRCA | Chemo + Pembrolizumab | 6 | PD | Lymph node | 1 |
| **RH18** | CRC | Atezolizumab + CD3 agonist | 2 | PD | Liver[1] | 2 |
| **RH19** | OV | Atezolizumab + BET inhibitor | 8 | SD | Lymph node | 3 |
| **RH21** | CRC | Atezolizumab + CD40 | 4 | PD | Liver[1] | 4 |
| **RH22** | HCC | Nivolumab + LAG3 antibody | 12 | CR | Liver[1] | 4 |
| **RH24** | GAC | Atezolizumab + CD40 | 4 | PD | Liver[1] | 2 |
| **RH25** | UPT | Atezolizumab + CD3 | 12 | PR | Liver[1] | 1 |
| **RH27** | BRCA | Atezolizumab + BET inhibitor | 2 | PD | Lymph node | 1 |
| **RH29** | OV | Atezolizumab + BET inhibitor | 4 | PD | Subcutaneous | 5 |
| **RH30** | MPM | Pembrolizumab | 1 | PD | Lymph node | 3 |
| **RH31** | NSCLC | Nivolumab | 4 | PD | Liver[1] | 2 |
| **RH33** | NSCLC | Pembrolizumab | 3 | PD | Lymph node | 0 |
| **RH34** | MPM | Atezolizumab + CD40 agonist | 20 | SD | Pleura | 3 |
| **RH35** | NSCLC | Pembrolizumab | 2 | PD | Liver[1] | 2 |

**Supplementary Table 2: Number of NARTs per screen.** Samples used for screening with the corresponding number of detected NARTs responses for each patient.

| | TIL | | REP TIL | | PBMC | | | |
|---|---|---|---|---|---|---|---|---|
| | **Before** | **After** | **Before** | **After** | **Before** | **After 1** | **After 2** | **Number of samples** |
| **RH08** | - | 0 | - | - | 3 | 1 | 2 | 4 |
| **RH10** | 0 | 0 | 0 | - | 0 | 0 | 0 | 7 |
| **RH11** | - | - | - | 8 | 8 | 5 | 1 | 4 |
| **RH13** | - | 0 | - | - | 0 | 0 | 1 | 4 |
| **RH15** | 0 | 0 | 0 | 0 | 0 | 0 | 0 | 7 |
| **RH16** | 0 | 0 | 1 | 1 | 2 | 1 | 1 | 7 |
| **RH17** | - | 0 | - | 0 | 0 | 0 | 0 | 5 |
| **RH18** | - | - | - | - | - | 0 | - | 1 |
| **RH19** | - | 0 | 0 | 0 | 1 | 0 | 1 | 7 |
| **RH21** | 0 | - | - | - | 2 | - | 0 | 3 |
| **RH22** | 0 | - | 0 | - | 0 | 0 | - | 4 |
| **RH24** | 0 | 0 | 0 | 0 | 2 | 1 | 4 | 7 |
| **RH25** | 0 | 0 | 0 | 0 | 1 | 2 | 5 | 7 |
| **RH27** | - | - | 0 | 0 | 2 | 2 | - | 4 |
| **RH29** | 0 | - | - | 0 | 0 | 2 | 0 | 5 |
| **RH30** | 0 | 0 | 0 | - | 0 | 0 | - | 5 |
| **RH31** | 0 | 0 | - | - | 8 | 6 | - | 4 |
| **RH33** | 0 | 0 | 0 | 0 | 14 | 8 | - | 6 |
| **RH34** | 0 | 0 | 0 | 0 | 3 | 0 | 3 | 6 |
| **RH35** | 0 | 0 | 0 | 0 | 2 | 1 | - | 6 |

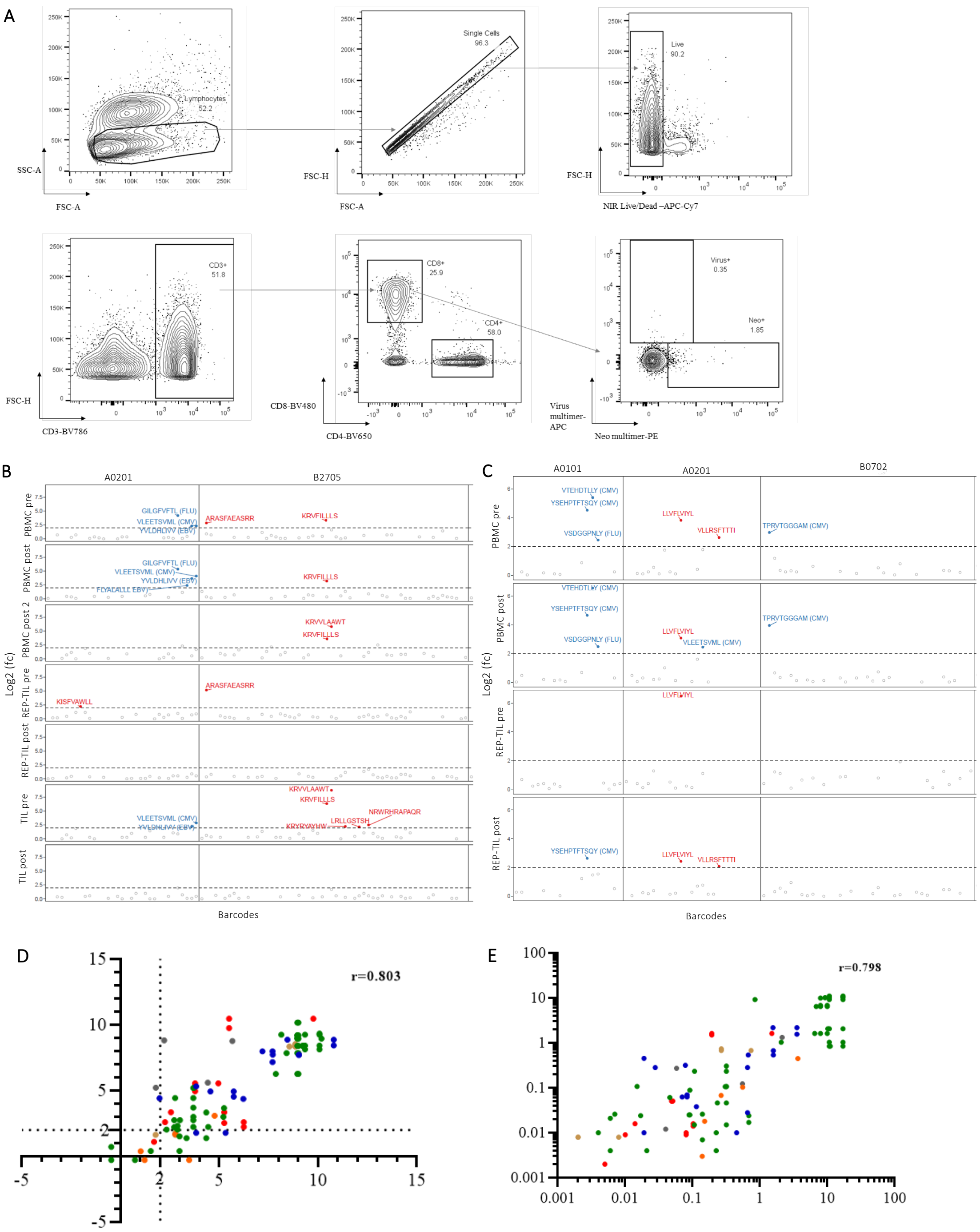


**Supplementary Figure 1. NARTs detection in patient samples. A)** Representative gating strategy for sorting multimer$^{+}$ CD8 T cells from bulk PBMCs. **B)** Summary plot for detected T cell reactivity for patient 25 (RH25). This plot is segregated vertically by the different PBMC and TIL samples, and horizontally by the HLA types included in the patient-specific panel. The Log2 fold change (fc) depicts DNA barcodes that have been positively enriched in the given sample, and a threshold Log2 (fc) value of > 2 depicts where significant T cell reactivity ($p < 0.001$) has been detected towards the neopeptide (red) or viral control peptide (blue). **C) ,** Summary plot for detected T cell reactivity for patient 27 (RH27). This plot is segregated vertically by the different PBMC and TIL samples, and horizontally by the HLA types included in the patient-specific panel. The Log2 fold change (fc) depicts DNA barcodes that have been positively enriched in the given sample, and a threshold Log2 (fc) value of > 2 depicts where significant T cell reactivity ($p < 0.001$) has been detected towards the neopeptide (red) or viral control peptide (blue). **D+E)** Correlation plots for detected T cell reactivity against viral epitopes in the healthy donor controls that were screened multiple times, represented as Log2 (fc) values (**D**) or estimated frequency values (**E**). The color scheme depicts the various healthy donor controls.

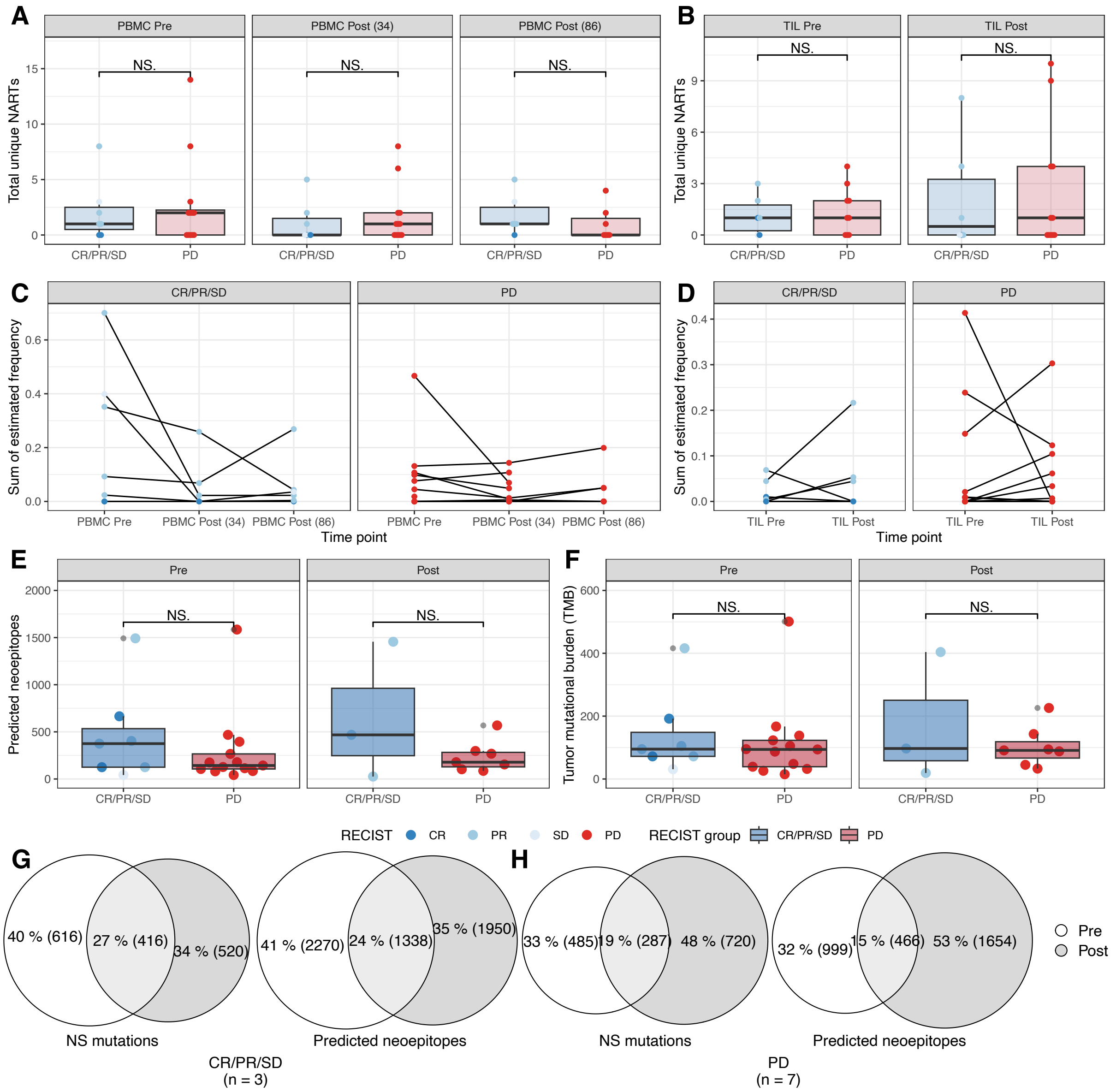


**Supplementary Figure 2. Tumor mutational burden and predicted neopeptides. A+B)** Total number of unique NARTs per patient pre- and post-treatment **(A)** in PBMCs for pre (p = 1), 34 weeks post-treatment (p= 0.59), and 86 weeks post-treatment (p= 0.30). In (B) in TIL pre-treatment (p = 0.81) and post-treatment (p = 0.83). **C+D)** Dynamic of the sum of estimated frequency of NARTs over time separated by patient outcome in (C) PBMC and (D) TIL. **E)** The number of predicted neoepitopes per patient in pre- and post-treatment separated by the patient outcome, pre (p=0.45) and post (p = 0.67). **F)** The tumor mutational burden per patient pre- and post-treatment separated patient outcome, pre (p= 0.63) and post (p =0.83). **G+H)** Venn diagram showing the overlap between the mutations and predicted neoepitopes pre- and post-treatment (G) for the CR/PR/SD and (H) for the PD group.

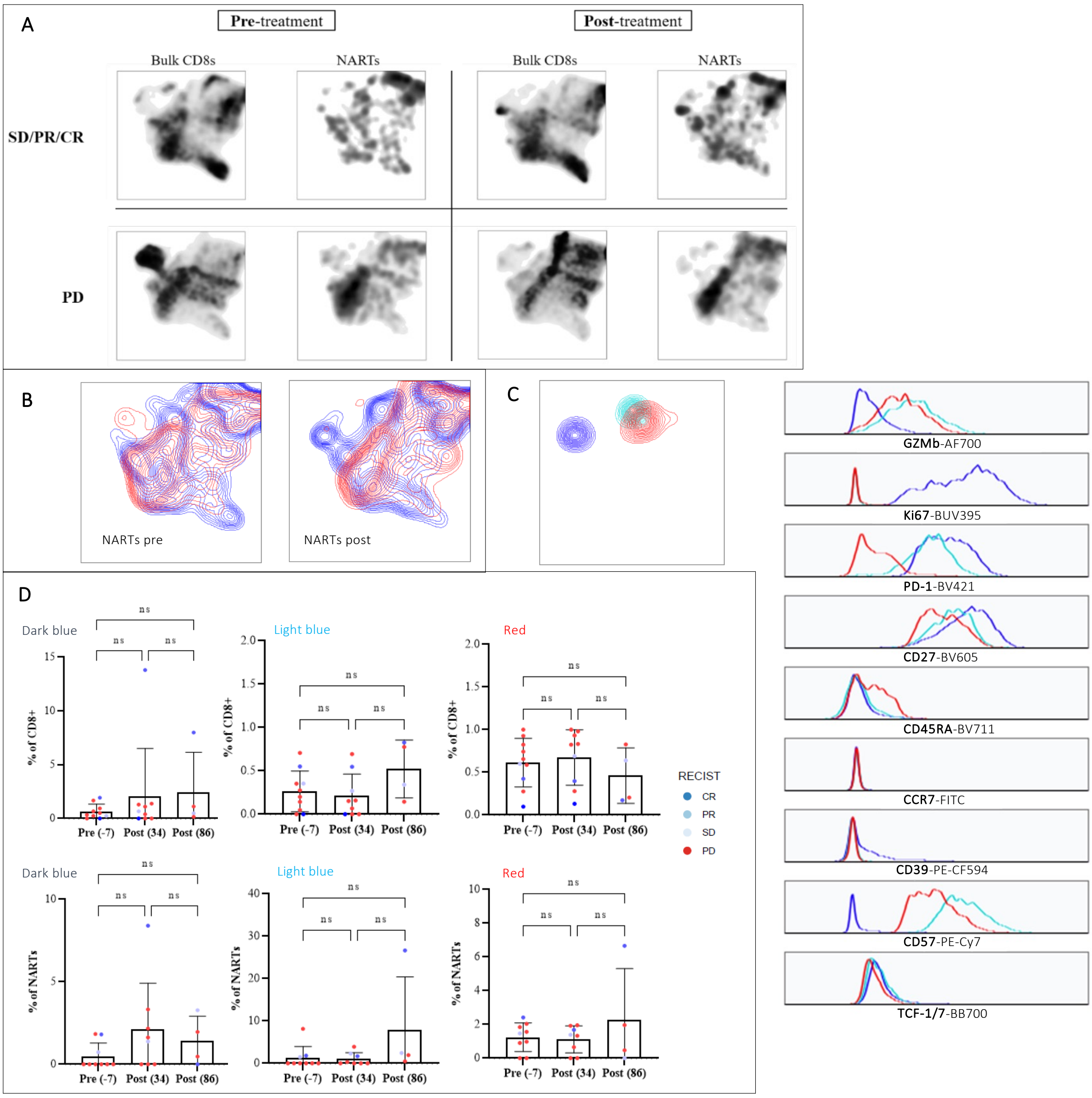


**Supplementary Figure 3, Phenotypic analysis of bulk CD8$^+$ T cells. A)** UMAP representation of the global structure of bulk CD8$^+$ T cells. NARTs for responding (top) and non-responding (bottom) patients, pre- and post-treatment, depicted as density plots. **B)** UMAP overlay of responding patient NARTs (blue) and non-responding patient NARTs (red), pre-treatment (left) and post-treatment (right). **C)** Selected NART sub-populations from the FlowSOM clustering and histograms for the corresponding phenotypic profiles. **D)** Quantitative assessment of the FlowSOM clustering for the selected sub-populations, depicting the frequency of CD8$^+$ T cells or NARTs that the particular sub-populations represent. For all the plots related to frequency of bulk CD8$^+$ T cells: (-7, n = 10; 34, n = 9; 86, n = 4) and for all plots related to frequency of NARTs: (-7, n = 9; 34, n = 8; 86, n = 4). CR; Complete responder, PR; Partial responder, SD; Stable disease, PD; Progressive disease. Significance for H is denoted based on Kruskal-Wallis test with Dunn's correction.
